# Identification, Isolation, Propagation and Inactivation of SARS-CoV2 Isolated from Egypt

**DOI:** 10.1101/2020.11.04.368431

**Authors:** M.G Seadawy, A.F Gad, M.F Elhoseny, B.El ELharty, M.D Shamel

**Author notes:** **Corresponding author**, <u>Mohamed Gomaa Seadawy, Mobile</u> No.

## Abstract

Severe Acute Respiratory Syndrome Coronavirus 2 causes the novel pandemic Pneumonia disease. It is a positive single strand ssRNA virus that infect human. COVID-19 appeared in Egypt in Feb 2020. The samples were taken from patients with COVID-19 symptoms at military hospital in Egypt and transported to the main chemical laboratories under all the biosafety measures according to WHO guidelines. All samples were tested with RT-PCR. Positive samples were cultured using VeroE6 cell lines. The propagated virus was isolated and inactivated. The isolated virus was sequenced using next generation sequencing and submitted into gene bank. This study provides an isolation, propagation and inactivation methodology which is valuable for production of inactivated vaccines against SARS-CoV2 in Egypt.

## Introduction

Severe acute respiratory syndrome coronavirus2 caused a pandemic disease called COVID-19. The COVID-19 appeared in China in 2019(**Zu. et al, 2019**). Egypt become infected on Feb 2020 with SARS-CoV2(**Abd El Dayem, Waleed A. 2020**). The pandemic virus consists of a positive single strand ssRNA. The spike protein of the virus is responsible for virus attachment with cell receptor angiotensin converting enzyme two (ACE2). The genome of SARS-CoV2 consists of two major portions(**Ana S., et al 2020**). The first is the genes responsible for replication ORF1a and ORF2b. The second portion is the genes coded for structural proteins like membrane, nucleocapsid, spike and envelope proteins. We identified the samples as positive SARS-CoV2 with RT-PCR. We used VeroE6 as cell line to propagate and isolate the virus. This study provides Identification, Isolation and Propagation methodology that enable researchers in Egypt to propagate SARS-CoV2 in high-titer for vaccine development and anti-viral drug screening.

## Materials and Methods

### Sampling

Oropheringial swabs were taken from symptomatic patients with COVID-19 from military hospital. The samples transferred, under full biosafety measures, in transportation media (sterile DMEM purchased from sigma Aldrich cat no. with ampicillin100IU/ml, streptomycin 100μg/ml) at 4 °C in biosafety transporting device to main chemical laboratories at ALMAZA-Cairo.

### Identification using RT-PCR

The RNA of all samples extracted using QIAamp viral RNA mini kit (QIAGEN, Hilden, Germany) following the manufacturer’s instructions. All samples were handled under a biosafety cabinet(germfree biosafety cabinet, SEA-III, 316 ss) according to laboratory biosafety guidelines main chemical laboratories in chemical warfare department, Egyptain army.

Molecular Identification was done using SARS-CoV-2 Real Time PCR detection KIT high profile. (Cat # VS-NCO212H) according to the manufacture instructions. The master mix was rehydrated with 15μL of rehydration Buffer and 5 μL of RNA of thermally treated and untreated samples were added (samples were run in triplicates), positive and negative controls (provided with the kit) were included in each test to judge the quality of amplification. Real time RT-PCR was done using Ariamx thermal cycler (Agilent, Germany) with the following parameters, reverse transcription step at 45oC/15 minute, followed by initial denaturing and enzyme activation step at 95oC/2 minute, then 45 cycled of denaturing at 95°C/10 sec and annealing/extension 60 °C/50 seconds with florescence collected at the end of this step.

### Propagation and Isolation

All the positive samples were were diluted with viral transfer medium containing Oropheringial swabs and antibiotics (Nystadin, penicillin-streptomycin 1:1 dilution) at 1:4 ratio and incubated for 1 hour at 4°C, before being inoculated onto VeroE6. Inoculated VeroE6 cells were cultured at 37°C, 5% CO2 in 1× Dulbecco’s modified Eagle’s medium (DMEM) supplemented with 2% fetal bovine serum and penicillin-streptomycin. Virus replication and isolation were confirmed through cytopathic effects and RT-PCR. Viral culture of SARS-CoV-2 was conducted in a biosafety Level-3 facility according to laboratory biosafety guidelines (germfree biosafety cabinet, SEA-III, 316 ss) according to laboratory biosafety guidelines of main chemical laboratories in chemical warfare department, Egyptian army.

### Inactivation

The Infected VeroE6 cells were scraped from the flask then centrifuged at 1000 rpm for 10 minute. The cell pellet was rinsed with 0.1M phosphate buffer saline (Thermo Fisher Scientific, USA) centrifuged at 1000 rpm for 10 min. Pellet was fixed 2.5% glutaraldehyde in 0.1 M Phosphate Buffer Saline at pH 7.4 for at least two hours at room temperature. Virus solution was added to Amicon Ultra 50,000 KDa NMWL (Cat No. UFC9003) and centrifuge at 4,000xg for 15 minutes to wash from excess glutaraldehyde. The sample was diluted with 1ml of PBS and centrifuge for 15 minutes at 4,000xg. The sample was diluted with 1ml of PBS and centrifuge for 3 minutes at 4,000xg. The concentrator was transferred to a clean tube and centrifuge at 1,000xg for 2 minutes to collect the concentrated sample. The sample was diluted in serum physiologic.

## Results

### Molecular Identification with Real Time PCR

Samples used in the current study were collected from clinically morbid patients with the classical symptoms of SARS-CoV-2 including hyperthermia (>38.5°C) dyspnea, dry cough, and atypical pneumonia as confirmed by chest X rays. Oropheringial swabs were taken and were confirmed positive for the presence of the virus using qRT-PCR. All before inoculation samples gave positive Cq value ranging from (21.26-26.51) for orf1 gene and (22.10-28.52) for the N gene. All after inoculation samples gave positive Cq value ranging from (14.52-17.31) for orf1 gene and (15.31-18.32) for the N gene (chart 1 &2).

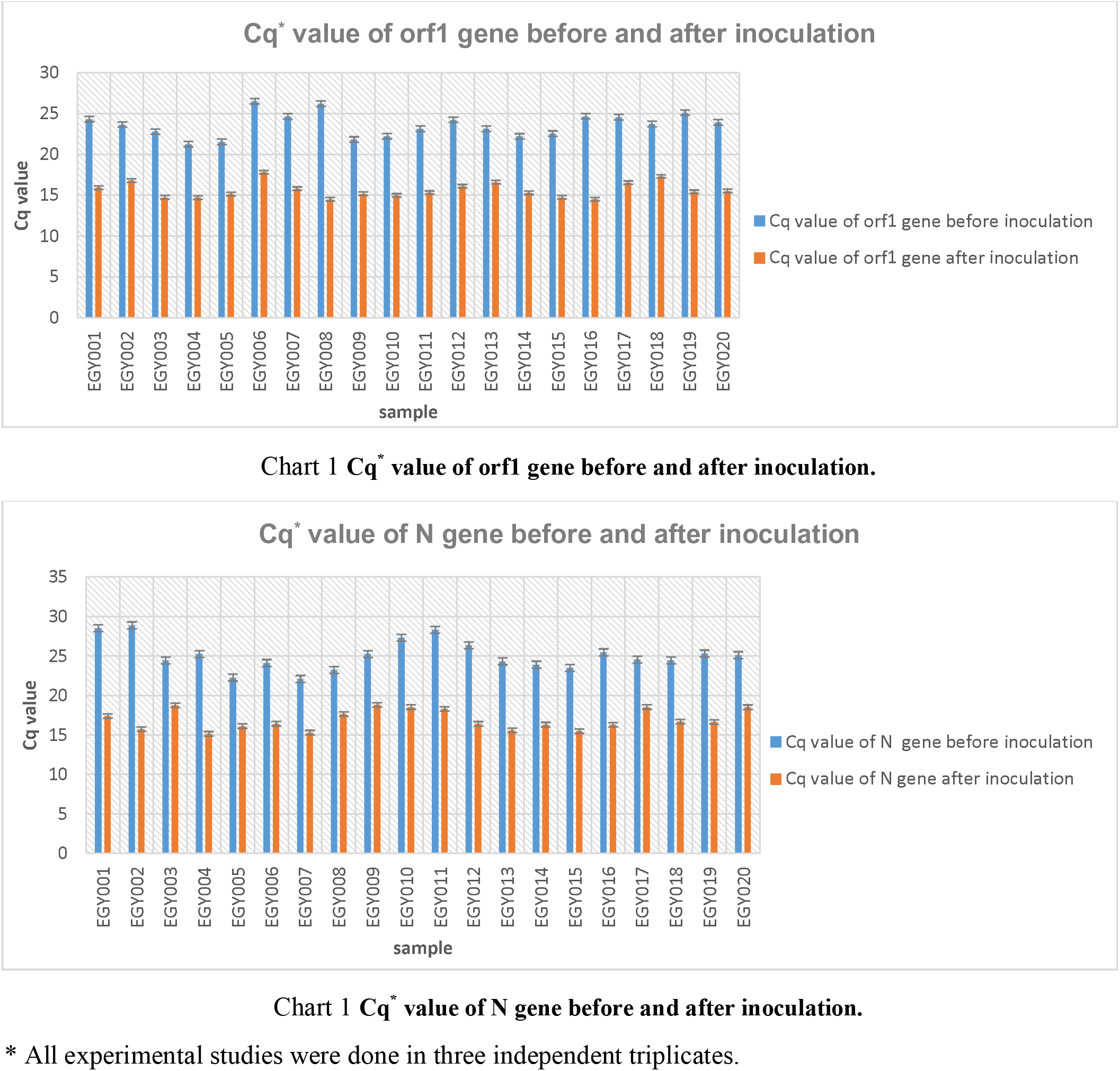

### Propagation of SARS-CoV2 on VeroE6 cell line

The cytopathic effect (CPE) of the SARS-CoV2 on the cells was followed and observed from the day 1 to day 4 after inoculation. Increasing cytopathic effects were recorded (Fig no 1). We wanted to identify the propagation of the virus as SARS-CoV-2 using RT PCR along with SARS-CoV-2 N gene and orf1 gene. The presence of SARS-CoV-2 in the propagated samples was proved in table 1 as we mention before. After a complete molecular identification of SARS-CoV2 with real time PCR (chart 1 &2), we sequenced a full genome of the virus for some sample and submitted into gene bank (accession numbers MT776904, MT798592, MT897260 and MT897261).

**Fig no 1.**
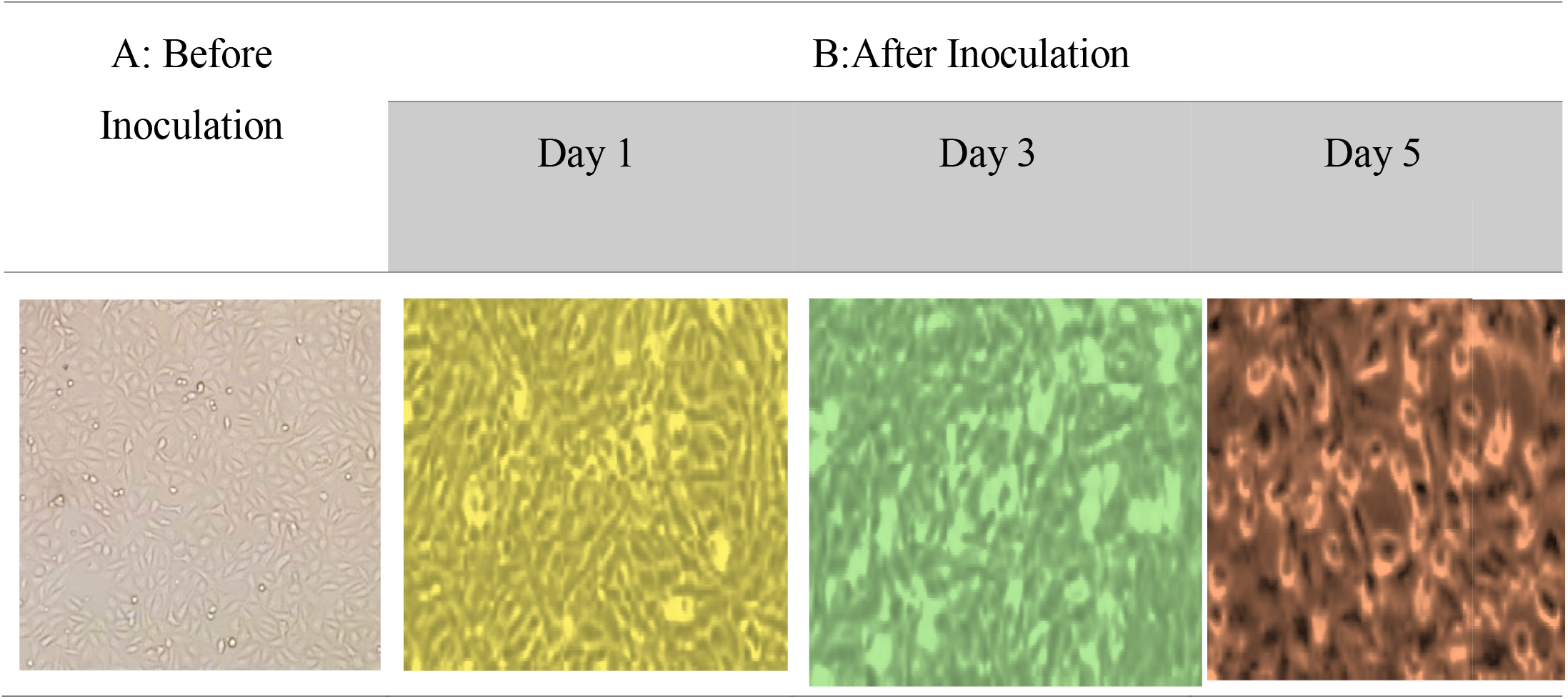
A: the cell line (VeroE6) before inoculation with SARS-CoV2. B: the cell line (VeroE6) after inoculation with SARS-CoV2 show gradually increasing the CPE at Day 1, Day 3 and Day 5.

### Inactivation

After inactivation of the propagated SARS-CoV2 in cell line as we mentioned before in method, we wanted to evaluate this inactivation process. Therefore, we inoculated the inactivated SARS-CoV2 again on VeroE6 cell line and did not observe any CPE.

## Discussion and Conclusion

SARS-CoV2 are enveloped ssRNA viruses that were distributed among humans in all over the world in 2019(**Ahmadi, et al. 2020**). This infection cause respiratory, enteric and neurological diseases (**Fang, et al.2020**). The emergence of coronaviruses at regular intervals poses an important threat to human health and economy of countries. This emphasizes the urgent need to develop effective vaccines to prevent future outbreaks and therefore isolation and inactivation of the SARS-CoV2 viruses become important issues.

In the process of specimen collection from patient and transfer to the laboratory, the process should be started and virus cultured immediately because lots of virus became inactivated. Therefore, we use transfer solutions, infection medium used in propagation process, to safe infectious SARS-CoV-2 during the transportation. We identified the SARS-CoV2 with real time PCR depending on orf1 and N genes detection according to RT-PCR kit. In addition, before inoculation the virus on the cells, we wash the cells with FBS free media to remove excess of fetal bovine serum (FBS). Then, to propagate virus, cells were suspended with a virus media including 2% FBS. Afterwards, we observed cytopathic effects from day 1 to day 5 after inoculation. Inactivation of SARS-CoV2 was done by the process we mentioned before to inactivate the virus for being used as inactivated vaccine but after complete consideration of inactivation vaccine production criteria.

As a conclusion, this study can be referenced for researchers in Egypt who aimed to propagate and isolate SARS-CoV-2 to work in vaccine production, genomics, sequencing, proteomics, gene editing like CRISPR and anti-viral drug screening.

## Data Availability

All data are available upon request from the corresponding author.

